# Profiling RT-LAMP tolerance of sequence variation for SARS-CoV-2 RNA detection

**DOI:** 10.1101/2021.10.25.465706

**Authors:** Esta Tamanaha, Yinhua Zhang, Nathan A. Tanner

## Abstract

The ongoing SARS-CoV-2 pandemic has necessitated a dramatic increase in our ability to conduct molecular diagnostic tests, as accurate detection of the virus is critical in preventing its spread. However, SARS-CoV-2 variants continue to emerge, with each new variant potentially affecting widely-used nucleic acid amplification diagnostic tests. RT-LAMP has emerged as a quick, inexpensive diagnostic alternative to RT-qPCR, but has not been studied as thoroughly. Here we interrogate the effect of SARS-CoV-2 sequence mutations on RT-LAMP amplification, creating 572 single point mutation “variants” covering every position of the LAMP primers in 3 SARS-CoV-2 assays and analyzing their effects with over 4,500 RT-LAMP reactions. Remarkably, we observed only minimal effects on amplification speed and no effect on detection sensitivity, highlighting RT-LAMP as an extremely robust technique for viral RNA detection. Additionally, we describe the use of molecular beacons to sensitively identify variant RNA sequences. Together these data add to the growing body of knowledge on the utility of RT-LAMP and increase its potential to further our ability to conduct molecular diagnostic tests outside of the traditional clinical laboratory environment.

## Introduction

Molecular diagnostic techniques are able to provide definitive identification of infectious agents through specific detection of DNA or RNA sequences of interest. However, target pathogens, and particularly RNA viruses, naturally display mutations and changes in their genomic sequences that can impact the sensitivity and accuracy of the molecular diagnostic test when the mutations occur in the regions targeted by the oligonucleotide primers and/or probes. The ongoing SARS-CoV-2 pandemic has seen the emergence of numerous viral variants from different regions of the world, with prominent effects on detection using molecular assays. For example, the B.1.1.7 “alpha” variant features a 6-base deletion (removing 2 amino acids of the spike protein, Δ69–70) which causes a failure of the S Gene assay in the widely used TaqPath^®^ COVID-19 multiplex RT-qPCR test (1). The other targets in this assay are unaffected, resulting in the ability to provide preliminary identification of variant RNA during detection, identifying a sample for further sequencing analysis and variant determination. In this particular case the other targets can be used for diagnostic detection, but the sensitivity of RT-qPCR to variant mutations is a significant concern to diagnostic testing given the worldwide reliance on the method. Early in 2021 the FDA issued guidance to molecular test developers (2) calling for evaluation of variant sequences in development of any molecular test, emphasizing the significance of understanding assay performance in the presence of targeted region mutations.

While much is known about the effects of mutations in qPCR assays (3, 4), loop-mediated isothermal amplification (LAMP) diagnostic assays are much more recent technology and accordingly we have comparatively little information on mutation effects. LAMP assays have been designed specifically to target SNPs in a sequence by placing the mutation directly at the 3′ or 5′ terminal base of both the FIP and BIP primers (5), but this is a highly artificial scenario. Natural mutations would most frequently consist of single base changes along a priming region of the LAMP primers, or more rarely, multibase deletions. For example, the B.1.1.7 Alpha variant contains the 6-base SGTF Δ69-70 and 9-base Orf1a SGF 3675–3677 deletions, but also >18 characteristic single base change mutations, whereas the B.1.617.2 Delta variant contains >20 point mutations and 2 deletions different from those in Alpha (6). In this study, we consider the 18 oligonucleotides making up 3 SARS-CoV-2 RT-LAMP assays. Mapping the primers onto SARS-CoV-2 sequences from GISAID shows numerous locations of mutation in small numbers of isolates worldwide, but a few (see 4 described in Methods) at a significant prevalence >10% of sequences deposited from at least 2 geographic locations independent from any of the well-known variants of interest or concern. Larger deletions likely have more detrimental effects on amplification than single base changes, though single base changes located close to the 3′ ends of primers might impact more than those located near the 5′ ends. Understanding the effects of single-base mutations on the effect of an amplification assay is critical to monitoring its continued ability to detect the target of interest as that target may naturally change over time.

In addition to reliable and consistent detection of the target, there is additional value in assays that can specifically identify variants of concern. Sequencing of positive samples will remain the most powerful and sensitive method for variant identification, but molecular diagnostic assays can present a significantly faster and cheaper approach if appropriately designed. The widely-used TaqMan probes rely on specific hybridization to targeted sequences and may display greater sensitivity to mutations than when they occur in primer regions; the Δ69-70 deletion occurs in the TaqPath probe region and completely prevents detection of RNA with that deletion. The SARS-CoV-2 variant strains B.1.1.7 (alpha), B.1.351 (beta), P.1 (gamma), B.1.617.2 (delta) and C.37 (lambda) contain numerous and distinctive mutations, most prominently located in the spike protein (1). While only Alpha carries the Δ69-70 deletion, all but Delta carry a 9-base deletion at positions 3675–3677 in the Orf1a sequence (termed the “SGF” deletion). This relatively large deletion provides a reliable means for designing molecular diagnostic assays that can distinguish between strains differing at this sequence location (1).

RT-LAMP has played a role in point-of-care and fieldable diagnostics, but during the SARS-CoV-2 pandemic has emerged as a prominent and widely-used molecular diagnostic method (7). Isothermal amplification can be performed with much simpler instrumentation that PCR, and detection of RT-LAMP can be conducted directly by visual color change (8, 9), fluorescence (10), sequence-specific probe such as DARQ (11) and molecular beacons (12), or coupled to secondary molecular analysis platforms such as CRISPR (13, 14) and next generation sequencing (e.g. LamPORE and LAMP-Seq)(15, 16). Several SARS-CoV-2 diagnostic protocols based on RT-LAMP are currently being used for large-scale test(13, 14, 17, 18) and can utilize crude or unpurified samples, saving time and cost while increasing testing flexibility and portability, exemplified by the utilization of RT-LAMP in the first molecular diagnostic for at-home use granted EUA in November 2020 (18).

As with qPCR, LAMP relies on oligonucleotide primers and mutations in the targeted regions may affect amplification efficiency. However, there is significantly more complexity in LAMP, as it utilizes 6 primers derived from 8 regions in the target sequence, with each of these regions similar in length to a PCR primer (19, 20). These 6 primers perform different roles during the amplification: F3 and B3 are located outside of the targeted region to aid in release of initial amplicons via strand displacement and are not incorporated into the amplification products. FIP and BIP are composed of two distinct target sequences and are the core primers responsible for the generation of the LAMP hairpin “dumbbell structure” and subsequent exponential amplification. LoopF and LoopB initiate from regions formed by “looping” from FIP and BIP and serve to further increase the speed of amplification.

And while previous studies of LAMP for SNP detection have described placing the mutation at the 3′ or 5′ terminus of both the FIP and BIP primers, this is very difficult to design in practice (5). A more general understanding of the impact of mutations on the 6 LAMP primers has not been characterized in detail. We set out to systematically investigate the effect of mutations by introducing single base changes at every position of each LAMP primer in 3 commonly used SARS-CoV-2 LAMP assays and running RT-LAMP with different concentrations of SARS-CoV-2 RNA to analyze the effects on assay speed and sensitivity. As RT-LAMP becomes a more widely-used method these results help us to assess the influence of mutations that may fall in the various primer regions.

In order to provide solutions for distinguishing sequence variants rather than just tolerating them, we also designed a variant-specific LAMP detection method based on molecular beacons. The use of molecular beacons containing locked nucleic acid has been previously demonstrated with LAMP (21, 22) and for SARS-CoV-2 detection (12) with beacons targeting the Gene S Δ69–70 deletion in the B.1.1.7 alpha variant (23). Here we designed new molecular beacons targeting the 9-base SGF deletion in Orf1a, enabling identification of multiple variants of concern including B.1.1.7, B.1.351, and P.1.

## Materials and Methods

### Single point mutation LAMP primers

Three previously described SARS-CoV-2 LAMP primer sets for SARS-CoV-2 were chosen to profile mutational position effects: As1e (24), E1, and N2 (9) (Table 1). Point mutations were identified in GISAID sequences as monitored by NEB Primer Monitor (primer-montor.neb.com), and 4 mutations reported in >10% of deposited sequences from >1 reporting location were chosen for LAMP screening. These mutations were: C2395T, in the As1e BIP primer; G2416T, in the As1e LoopB primer; T29148C, in the N2 F3 primer; and G29179T, in the N2 LoopF primer (locations seen noted in Table 1 below). We then modeled these produced mismatches (G:T, C:T, T:G, G:A) by introducing a mismatch at every base position for each of the primers, changing: C➔T, T➔C, G➔T, or A➔C. The resulting 572 primers were synthesized in 96-well plates at 10x concentration (2 μM F3, B3; 16 μM FIP, BIP; 4 μM LoopF, LoopB) by Integrated DNA Technologies (IDT) and spot-checked for concentration using 90 of the provided oligos.

**Table 1.**
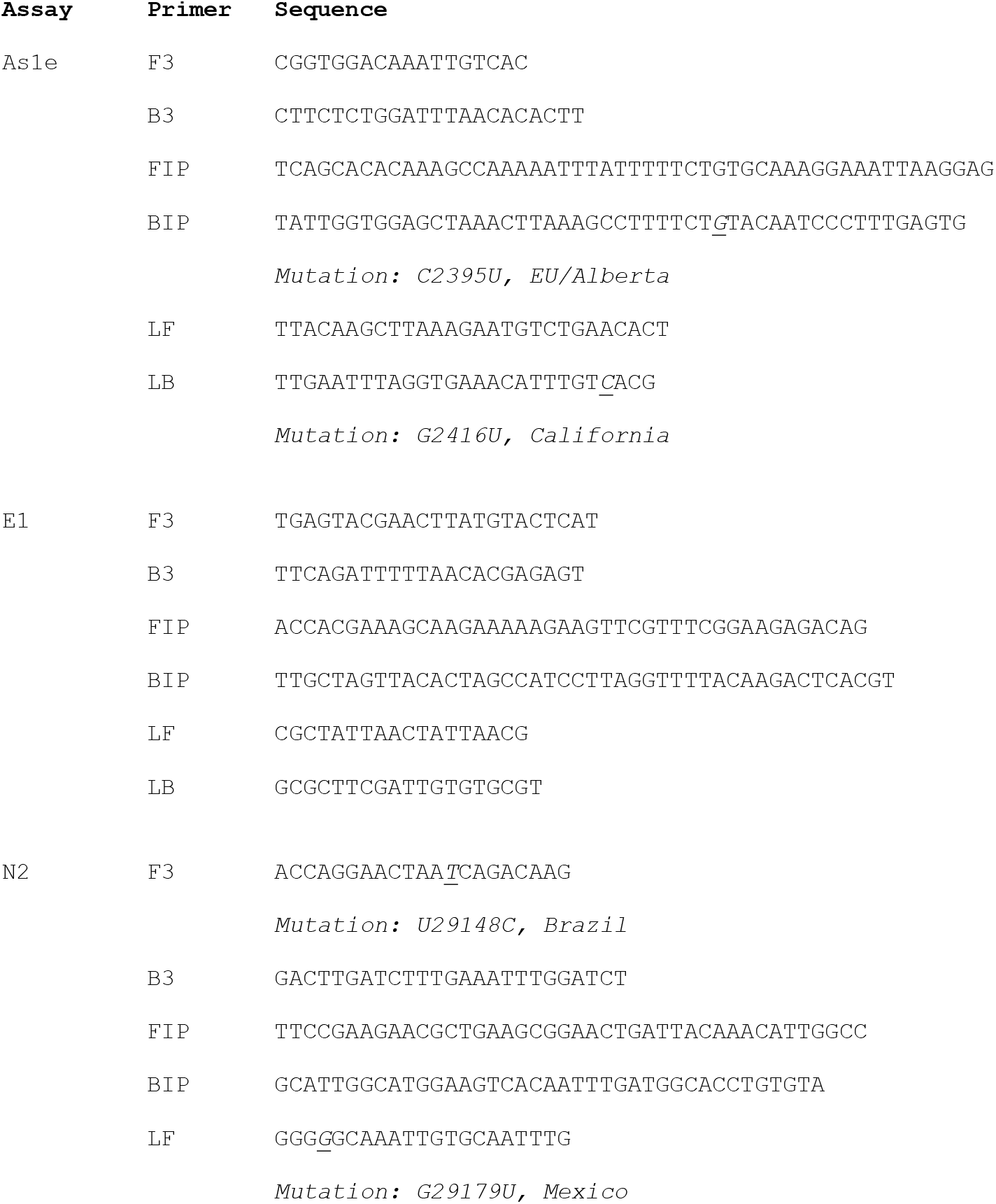
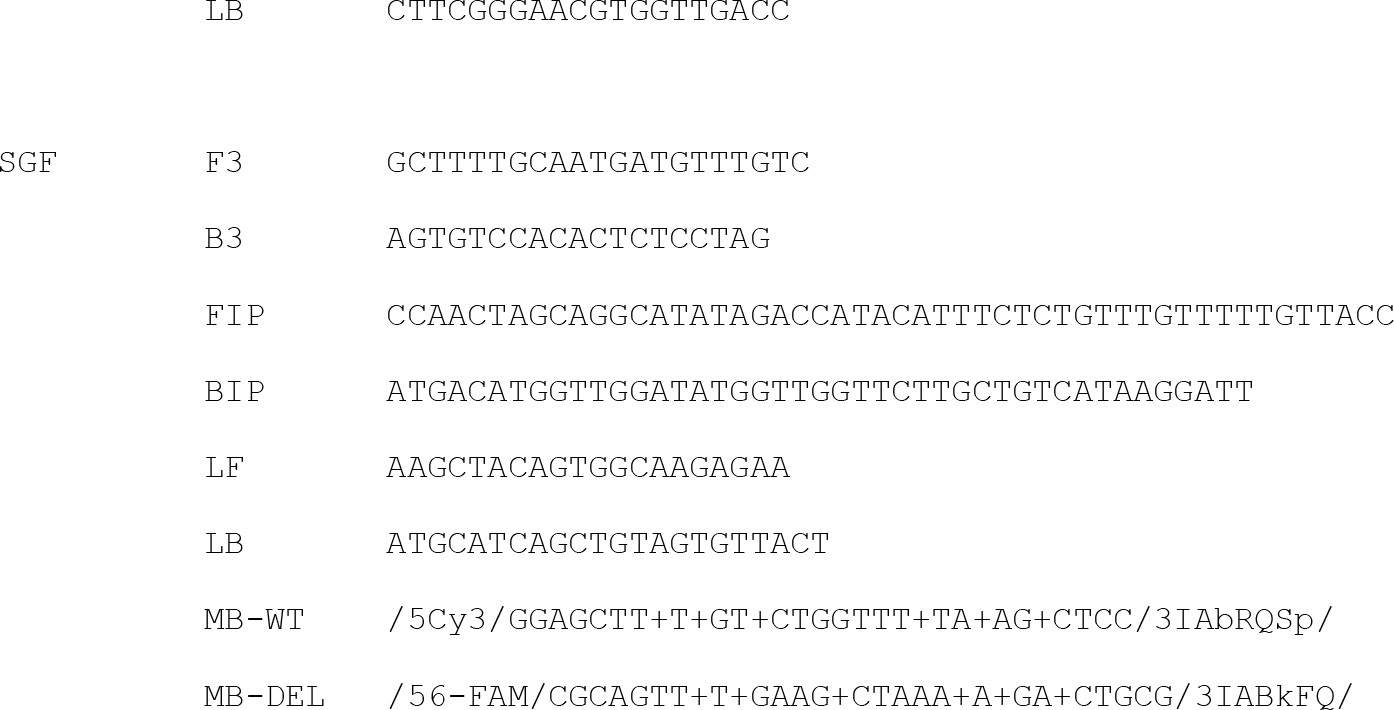
LAMP Primers

### LAMP primers and molecular beacons

LAMP primers were designed with the SGF deletion located between B1c and LB (Figure 1) using the NEB LAMP Primer Design Tool (https://lamp.neb.com) and with enough length between B1c and LB to accommodate the location of molecular beacons in this region. Molecular beacons targeting either wild-type or the SGF deletion sequence were designed using principles according to (12). As the deletion is located in a relatively AT-rich region, the annealing temperatures for these beacons are lower: the calculated Tm of the annealed beacon-target for wt MB and SGFdel MB is 63.9 and 62.5 °C, and the stem is 55.1 and 54.7 °C, respectively. These beacons were synthesized as Affinity Plus qPCR Probes by IDT with sequences shown in Table 1.

**Figure 1.**
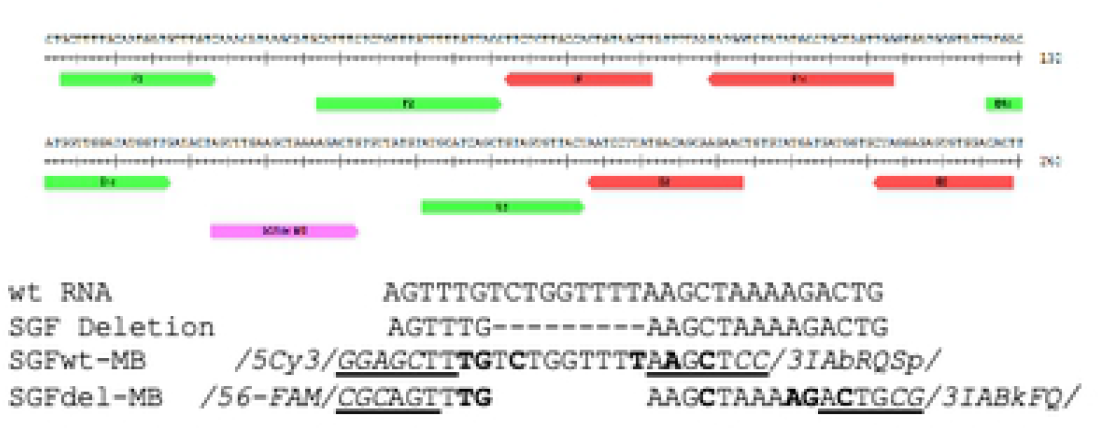
Design of SGF LAMP primers and beacons. Upper panel shows the locations of various LAMP primers and molecular beacons. The lower panel compares the sequences for wt, SGF deletion, SGFwt-MB and SGFdel-MB. Dashes: bases deleted in SGF deletion; Bold: LNA base; Underlined: stem region; Italics, non-target sequence, attached fluorophores and quenchers.

### RT-LAMP reactions

RT-LAMP reactions were performed using WarmStart^®^ LAMP Kit (DNA & RNA) (E1700) with standard primer concentrations (0.2 μM F3, 0.2 μM B3, 1.6 μM FIP, 1.6 μM BIP, 0.4 μM Loop F, 0.4 μM Loop B) in the presence of 40 mM guanidine hydrochloride (9) in 25 μl on 96-well plates at 65 °C in a Bio-Rad CFX96 instrument. LAMP amplification was measured by including 1X NEB LAMP Dye (B1700) or 1 μM SYTO™-9 (ThermoFisher S34854), 0.5 μM SGFdel or 1.0 μM wt beacon, and fluorescent signal was acquired at 15 second intervals. Synthetic SARS-CoV-2 RNAs were obtained from Twist Bioscience (Control 2 for WT MN908947.3; Control 14 for B.1.1.7; Control 16 for B.1.351; and Control 17 for P.1) and diluted in 10 ng/μl Jurkat total RNA based on the copy number provided by the manufacturer.

## Results

### Positional Mutation Effects

To mimic the effect of a potential SARS-CoV-2 variant in an RT-LAMP assay, we focused on single point mutations at each primer base position and the SGF deletion that is found in several variants of concern. For the single point mutation primers, each of the 527 variant primers from the 3 assays (As1e, N2, E1) was tested in RT-LAMP reactions with three different SARS-CoV-2 RNA copy number concentrations: 100, 200, and 10,000 copies in order to gain a sense of the mutation effect on reaction speed and sensitivity. Both lower concentrations allowed for amplification effects to be confidently determined outside of stochastic performance when close to the limit of detection (LOD ~50 copies) in the 100 copy reactions, particularly for the As1e primer set which displays slightly lower sensitivity in our testing. The reaction output speed was measured relative to the fully-complementary wild-type primer and plotted against the position in the primer sequence (Figure 2). Overall, the ~4,500 RT-LAMP assays containing single point mutations within any of the six primers resulted in minimal to no effect on the ability to amplify the target at any of the copy numbers used, regardless of primer and gene target. The most common result observed was a 5–10% reduction in amplification speed in the presence of a mismatch.

**Figure 2.**
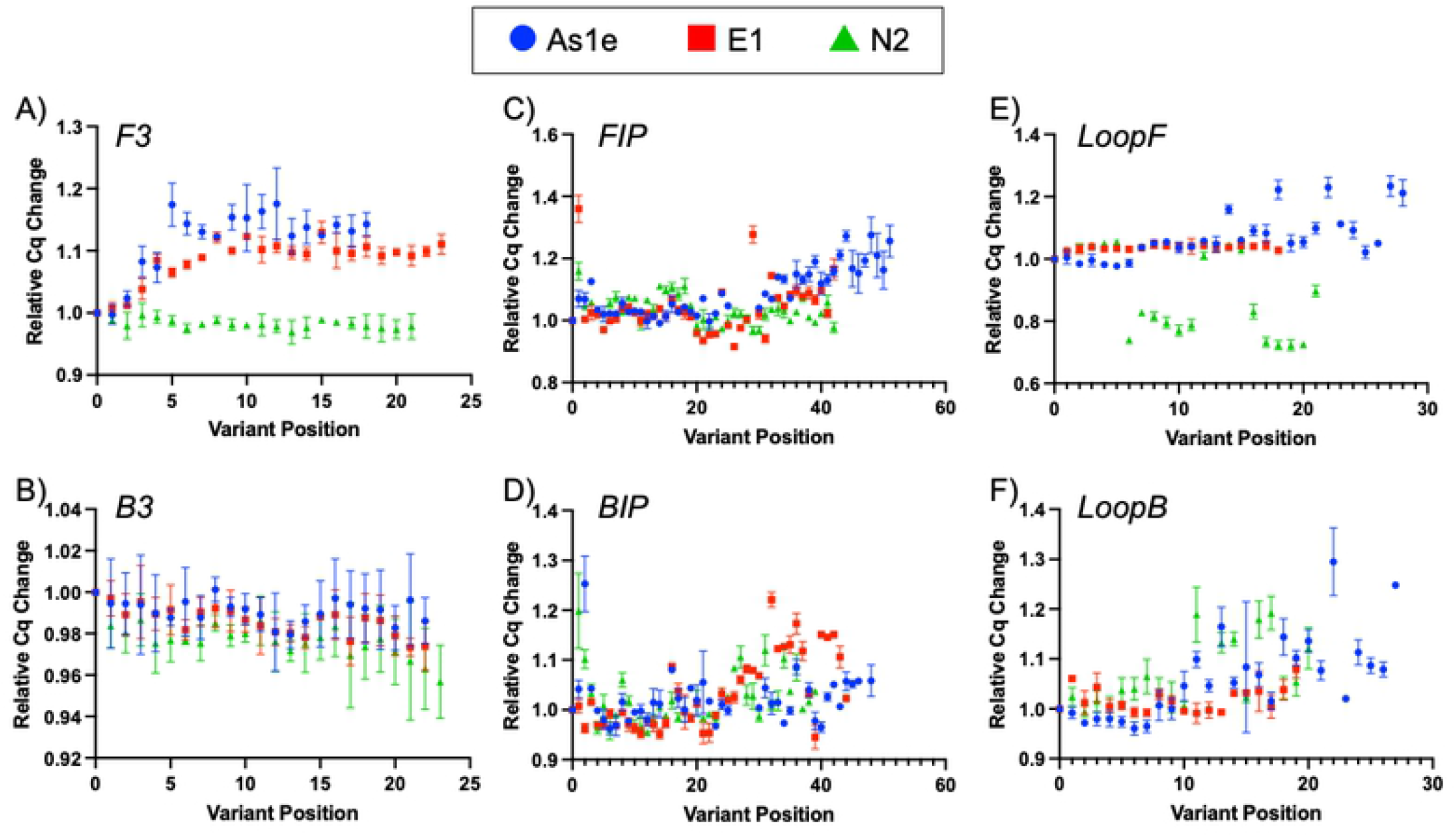
Mutation position effects on RT-LAMP amplification. Plots of the effects of change relative to the WT primer set for all three genes explored, As1e (blue circle), E1 (red square), N2 (green triangle). (A) F3 primer, (B) B3 primer, (C) FIP primer, (D) BIP primer, (E) Loop F primer, (F) Loop B primer.

Evaluating the effects across the different primers revealed some minor differences based on position and role of the primer. The B3 primers showed remarkably little impact in any of the 69 variant primer sets for the 3 amplicons, though the F3 primer did see some slowing when mismatches were present away from the 5′ end in the E1 and As1e sets (Fig. 2A, B). These primers are the least critical to the reaction, but the difference between the two may indicate differential mismatch tolerance during reaction initiation (B3 annealing to single-stranded RNA and extension by RTx reverse transcriptase) vs. the strand invasion and displacement via *Bst* 2.0 polymerase that occurs at the F3 primer. With the more critical FIP and BIP primers, the 3′ half of the primer (F2/B2) serves to invade and primer double-stranded DNA, with the 5′ half annealing to displaced product strands to form the ‘loop’ dumbbell shapes for amplification. As shown in Figure 2C-D more of the variant primer sets caused amplification delays relative to the fully base-paired control when the mutations were located toward the 3′ end of the FIP and BIP (F2/B2 regions) in all 3 LAMP assays. The extreme 5′ end displayed an increased mismatch effect on reaction speed, likely indicating an impact on polymerase extension from a mismatch in the looped-back LAMP hairpin structure, but detection sensitivity was not impacted even with those primer sets. Surprisingly the speed of the N2 assay increased when mutations were present at internal and 3′ positions of the LoopF primer. This was not a consistent effect, and while difficult to anticipate from sequence prediction its underlying mechanism will be investigated further in future LAMP assays. As a summary of the effects Table 2 lists the number of positions from each primer that resulted in a change of more than 10% in time to detection from the WT primer baseline. Though overall effects on amplification were minimal, the greater impact of 3′ mutations is clear from the trends in Figure 2. And while a significant number of variant primers resulted in decreased reaction speeds, in all 527 variant primers tested no mutation position prevented amplification in the RT-LAMP reaction with SARS-CoV-2 RNA even with lower RNA copy numbers.

**Table 2.** Effect of Single Point Mutations

| Primer | No. Positions with<br>>10% LAMP Time<br>Change | Total<br>Bases | Fraction<br>Positions<br>with >10%<br>Effect |
| --- | --- | --- | --- |
| F3 | 22 | 62 | 0.35 |
| B3 | 0 | 69 | 0 |
| FIP | 27 | 131 | 0.21 |
| BIP | 21 | 128 | 0.16 |
| LF | 18 | 68 | 0.26 |
| LB | 13 | 66 | 0.19 |

### Deletion Detection with Molecular Beacons

To investigate the effect of a deletion on an RT-LAMP assay, we designed a primer set targeting the SGF deletion region (Orf1a 3675–3677) with two molecular beacons targeting either the WT or the variant target region (Fig. 1). We initially evaluated this primer set for specificity and sensitivity and found it was able to detect both WT and variant RNAs with similar sensitivity of approximately 50 copies and with no apparent non-template amplification signal in 40 minute reactions (data not shown). We then evaluated detection using the molecular beacons with 10-fold dilutions of WT or B.1.1.7 synthetic RNA and compared them with conventional intercalating fluorescence detection. Both molecular beacons detected their intended targets as designed with robust specificity and even at 10,000 copies of target RNA they recognized only their intended amplification products (Fig. 3). This is in contrast to another strategy where the deletion was placed at the ends of the FIP and BIP (5) and detected with an intercalating dye. With this design, WT vs. B.1.1.7 discrimination was efficient at low (≤1,000 copies RNA) inputs but at higher copy numbers, the two variant sequences could only be distinguished by their amplification speed.

**Figure 3.**
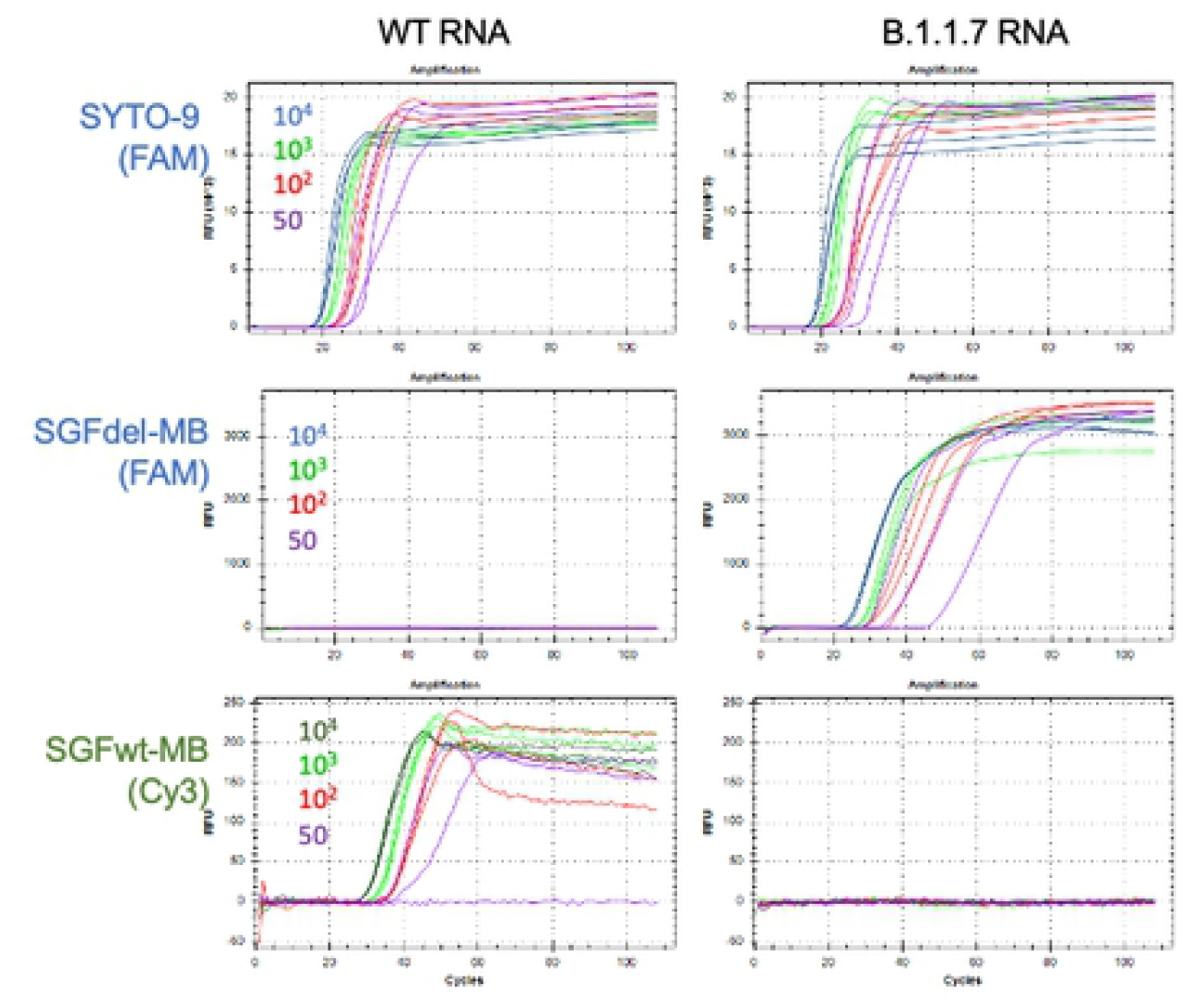
Detection of target RNAs by SGFwt and SGFdel molecular beacons. LAMP reactions with either WT RNA (left panels) or B.1.1.7 RNA (right) in the presence of SYTO-9, SGFdel-MB, or SGFwt-MB. The primer set amplifies both the wt and B.1.1.7 RNA with similar efficiency as detected with SYTO-9 (top). When beacon was added as a reporter, both SGFdel-MB (middle) and SGFwt-MB (lower) showed only with their intended template RNAs from 50–10,000 copies.

We then assessed the sensitivity of the SGFdel beacon to detect variant RNA containing the SGF deletion. We performed 24 repeats of LAMP reactions each with 50 copies of synthetic RNAs from B.1.1.7, B.1.351, or P.1 and the reactions were detected by either the molecular beacons or by dsDNA binding dye. The results indicated that both detection approaches were able to detect all the variant RNAs with similar efficiency (Table 3). We further tested combining both molecular beacons in the same LAMP reaction for identification of the input viral RNA could be determined by running 24 repeat reactions in the presence of both SGFwt and SGFdel molecular beacons. Results from these reactions showed that target RNAs were reliably detected with the same level of sensitivity when both beacons were present, and we observed no interference between the two beacons (Table 4).

**Table 3.** Specific Detection of Variant RNA with LAMP and Molecular Beacons

|  | *RNA* |  |  |  |
| --- | --- | --- | --- | --- |
|  | WT | B.1.1.7 | B.1.351 | P1 |
| SYTO-9 | 21 | 21 | 23 | 24 |
| Beacon | 18 | 17 | 24 | 23 |

*positives from 24 repeats, 50 copies/reaction*

**Table 4.**
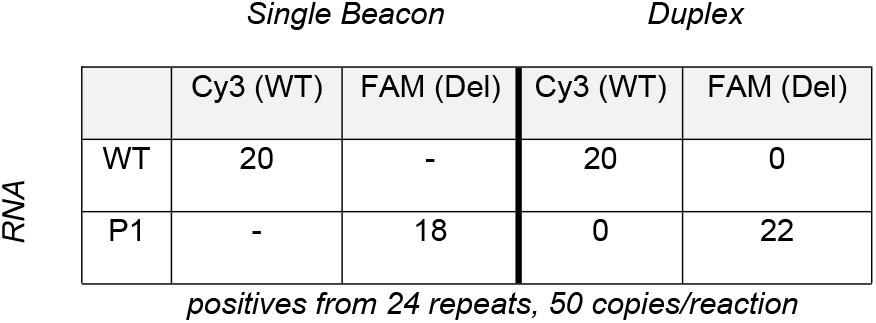
Dual-beacon RT-LAMP for Variant RNA Detection

## Discussion

RT-LAMP has become a significant molecular diagnostic tool during the COVID-19 pandemic due to its simplicity and flexibility, expanding the reach of molecular methods beyond the clinical laboratory where RT-qPCR remains the dominant method. Building a greater understanding of how RT-LAMP assays perform will be critical to increasing their utility, and tolerance to mutations is a pressing need for this and any future viral detection effort. Here we established the first comprehensive screen of LAMP primer tolerance to mutation, investigating a single base mutation at every position of every primer in three prominent SARS-CoV-2 RT-LAMP assays.

Remarkably, we find very little impact of the single base changes, with only marginal effect on amplification speed in most positions. In comparison, single-base changes in SARS-CoV-2 mutations have been demonstrated to have significant impact on RT-qPCR assays. One N gene mutation (G29140U) falling in a Forward PCR primer resulted in a 5–6 C_q_ increase with an estimated 67-fold drop in assay sensitivity (25).

Another N gene single base change (G29179U) in the CDC N2 Forward primer produced a ~4 C_q_ increase in PCR and a ~5-fold decrease in sensitivity (26). Both of these fall at internal positions in the PCR primers but cause significant effects while equivalent positions in LAMP primers showed minimal impact. Effects were more pronounced near the 3′ end of the FIP and BIP primers indicating the importance of initiation *via* the F2 and B2 regions in driving the formation of the LAMP dumbbell structures. However, while the speed increases were more pronounced at those positions, effect was minimal as compared to equivalent mutation positioning in PCR, where a 3′ mismatch causes a >100-fold sensitivity effect and can be used to completely discriminate a SNP due to inefficient mismatch extension (3). The robustness of RT-LAMP to sequence variation is a significant benefit over RT-qPCR, with reduced worry about deleterious effects from the commonly emerging single-base changes that could occur with some frequency in the regions targeted by the LAMP primers. Additionally, many RT-LAMP assays combine primer sets for added speed and sensitivity (9), adding an additional layer of protection against possible sequence variation.

The converse of this assay robustness however is an inherent difficulty in identifying variants during the amplification reaction. While sequencing offers greater confidence and detail for variant calling the ability to utilize the diagnostic amplification for prospective variant identification as with the TaqPath S-gene dropout can be a valuable feature of a potential diagnostic method. We observed difficulty targeting even a large 9-base deletions by typical LAMP primer design alone, but by utilizing a molecular beacon approach as first described by (23) we were able to accurately amplify and identify RNA from the three SARS-CoV-2 variant sequences containing the SGF deletion. By combining the beacons for the wild-type and deletion sequence, we could call wild-type or variant based on the detected sequence, indicating the potential ability for variant calling in the RT-LAMP assay by multiplexed beacon design.

Taken together these data position RT-LAMP as an attractive diagnostic method with a high level of tolerance to common sequence mutations. Recent FDA guidance described a need for understanding this tolerance for any molecular diagnostic test, and use of RT-LAMP could convey increased confidence to developers that a test will maintain performance. In situations where variant identification is desired, the robustness of LAMP primers is a detriment, but use of molecular beacons provides a sensitive addition to specific sequence detection with fluorescence detection. RT-LAMP remains a promising molecular diagnostic method and by further characterizing its behaviors and increasing its applicability we hope to further enable it to bring diagnostic testing to field and point-of-care settings where its advantages can be more fully utilized.

## Acknowledgements

The authors thank Eric Hunt (NEB) for assistance designing the variant primer panels and Nicole Nichols (NEB) for inspiration to conduct the full set of variant profiling experiments. We are grateful to New England Biolabs for funding the work and fostering an environment of scientific discussion and collaboration, without which this work would not have been possible.

## Supporting Information

**Supporting Figure 1.** Mutation position effects on RT-LAMP amplification at 100 copies of SARS-CoV-2 RNA.

Plots of the effects of change relative to the WT primer set for all three genes explored, As1e (blue circle), E1 (red square), N2 (green triangle) at 100 copies of SARS-CoV-2 RNA. (A) F3 primer, (B) B3 primer, (C) FIP primer, (D) BIP primer, (E) Loop F primer, (F) Loop B primer

**Supporting Figure 2.** Mutation position effects on RT-LAMP amplification at 200 copies of SARS-CoV-2 RNA.

Plots of the effects of change relative to the WT primer set for all three genes explored, As1e (blue circle), E1 (red square), N2 (green triangle) at 200 copies of SARS-CoV-2 RNA. (A) F3 primer, (B) B3 primer, (C) FIP primer, (D) BIP primer, (E) Loop F primer, (F) Loop B primer

